# Habitat size, health and saturation do not alter movement decisions or the preference for familiarity in a social coral-reef fish

**DOI:** 10.1101/2021.06.17.448906

**Authors:** Catheline Y.M. Froehlich, Siobhan J. Heatwole, O. Selma Klanten, Marian Y.L. Wong

## Abstract

While habitat is often a limiting resource for group-living animals, we have yet to understand what aspects of habitat are particularly important for the maintenance of sociality. As anthropogenic disturbances rapidly degrade the quality of many habitats, site-attached animals are facing additional stressors that may alter the trade-offs of moving or remaining philopatric. Here we examined how **habitat health, size and saturation** affect movement decisions of a coral-dwelling goby, *Gobiodon quinquestrigatus*, that resides within bleaching-susceptible *Acropora* coral hosts. To assess effects of habitat health, we translocated individuals far from their home corals into dead corals with the choice of adjacent healthy corals. To assess effects of habitat size and saturation, we manipulated coral sizes and the number of residents in healthy corals. Remarkably, 55% of gobies returned home regardless of treatment, 7% stayed in the new coral, and the rest were not found. Contrary to expectations, habitat factors did not affect how costs of movement influence group-living decisions in this species. These site-attached fishes preferred to home instead of choosing alternative habitat, which suggests a surprising awareness of their ecological surroundings. However, disregarding alternative high-quality habitat is concerning as it may affect population persistence under conditions of rapid habitat degradation.

## Background

Social animals often live in specific microhabitats, like tunnels for mole rats, sponges for shrimp, and cnidarians for reef fishes [1–3]. For many social animals, such habitat provides access to food, mates, territory and breeding sites [4–6], and therefore represents a key limiting resource [1–3]. As such, habitat can play a key role in the evolution and maintenance of sociality, and habitat factors are known to modulate decisions of many taxa to remain in groups or move to breed elsewhere [7–11].

According to the ecological constraints hypothesis [8], delaying reproduction to remain in groups outweighs moving to other habitat to breed independently due to high costs of movement and habitat saturation [7,8,11,12]. Movement imposes substantial costs because of predation risk and energy expenditure, especially if alternative habitat is already saturated, which could arise from certain life history characteristics [13]. Alternatively, when reproduction of low ranking individuals is suppressed, moving to less saturated habitats could mean reaching breeding positions sooner [7,14]. Hence for social animals, the trade-offs between dispersing and remaining philopatric are likely driven by both habitat saturation and costs of movement. Alternatively, the benefits of philopatry hypothesis suggests that remaining in groups enables access to high quality habitat, which can increase survival and long-term reproduction [8,9]. Habitat quality is often inferred via habitat size, and larger habitats typically support larger groups due to the additional space and resources available for supporting more individuals and reducing conflict [15,5,16]. Lower ranking individuals may even forgo reproduction to reap the benefits of remaining in larger habitat [3,5].

While studies have focused primarily on the role of habitat size as a measure of quality [5,15,17,18], other parameters clearly dictate habitat quality and hence the degree of movement and sociality of animals. For social animals residing in living habitats, as is seen in shrimp inhabiting sponges [2], ants inhabiting plants [19], and fish inhabiting cnidarians [3], movement decisions may depend on the health of their ‘host’ habitat. Given that habitat degradation is occurring at an alarming rate due to environmental and anthropogenic disturbances [20,21], investigating the role of habitat health is necessary for a holistic understanding of how habitat promotes sociality [11]. Understanding the interaction between habitat health, size, and saturation on the movement and sociality of habitat-specialists is especially important since threats of habitat degradation and mortality are increasing. Therefore, we urgently need to assess the interplay between multiple habitat factors on movement decisions in order to predict and potentially mitigate the social consequences of environmental degradation.

Here we investigated how multiple ecological factors, namely habitat size, health, and saturation, influence sociality and movement decisions using a social coral-dwelling goby *Gobiodon quinquestrigatus* (Gobiidae) as our model species. Coral gobies provide an excellent model system since they reside within branches of living acroporid corals [22] and are site-attached even after coral bleaching [23]. *Gobiodon quinquestrigatus* are classified as facultatively social because group-living only occurs when coral hosts are large enough, whereas pair-forming occurs when corals are small [24]. Such facultative sociality is useful because it enables us to examine and manipulate the potential factors promoting group-over pair-formation. We completed an *in situ* manipulative experiment to test the predictions that gobies would prefer to move to: (1) healthier versus bleached/dead habitat, (2) larger habitat, and (3) habitats with smaller groups (less saturated) to improve breeding opportunities.

## Methods

### Site location

Experiments were completed on SCUBA during two trips (Sep-Nov 2018 and May-June 2019) at four inshore reefs near Mahonia Na Dari Research and Conservation Centre in Kimbe Bay, West New Britain, Papua New Guinea (−5.42896°, 150.09695°).

### Experimental design

We completed our study *in situ* by removing a goby from its home coral and translocating it into a dead coral that was situated adjacent to a live coral. To set up the experiments, dead corals of *Acropora kimbeensis* were opportunistically located on the reef. These dead corals were of two size categories: small (11.2-cm avg. diameter) and large (17.3-cm avg. diameter). We then randomly searched for similarly-sized live corals that contained *G. quinquestrigatus* individuals. To set up one trial, a dead coral was placed within 10 cm of the similarly-sized live coral (Fig 1, Suppl Fig 1a,b & 2). In neighbouring corals (within a 10-m radius), we then located a ‘focal’ *G. quinquestrigatus* individual that was smaller (16.9-mm avg. standard length, range: 12.2-22.5 mm) than gobies in the live coral (next to the dead coral). Selecting a smaller goby was important to reduce potential eviction by residents [18]. The focal goby was removed from its original home coral using a clove oil anaesthetic solution and hand nets [25] and injected with a unique visible implant elastomer identification tag (Northwest Marine Technology, Inc.) [22]. The focal goby was then translocated into the dead coral (Fig 1), and we revisited trials daily for up to 7 days to determine where the focal goby subsequently moved.

**Fig 1.**
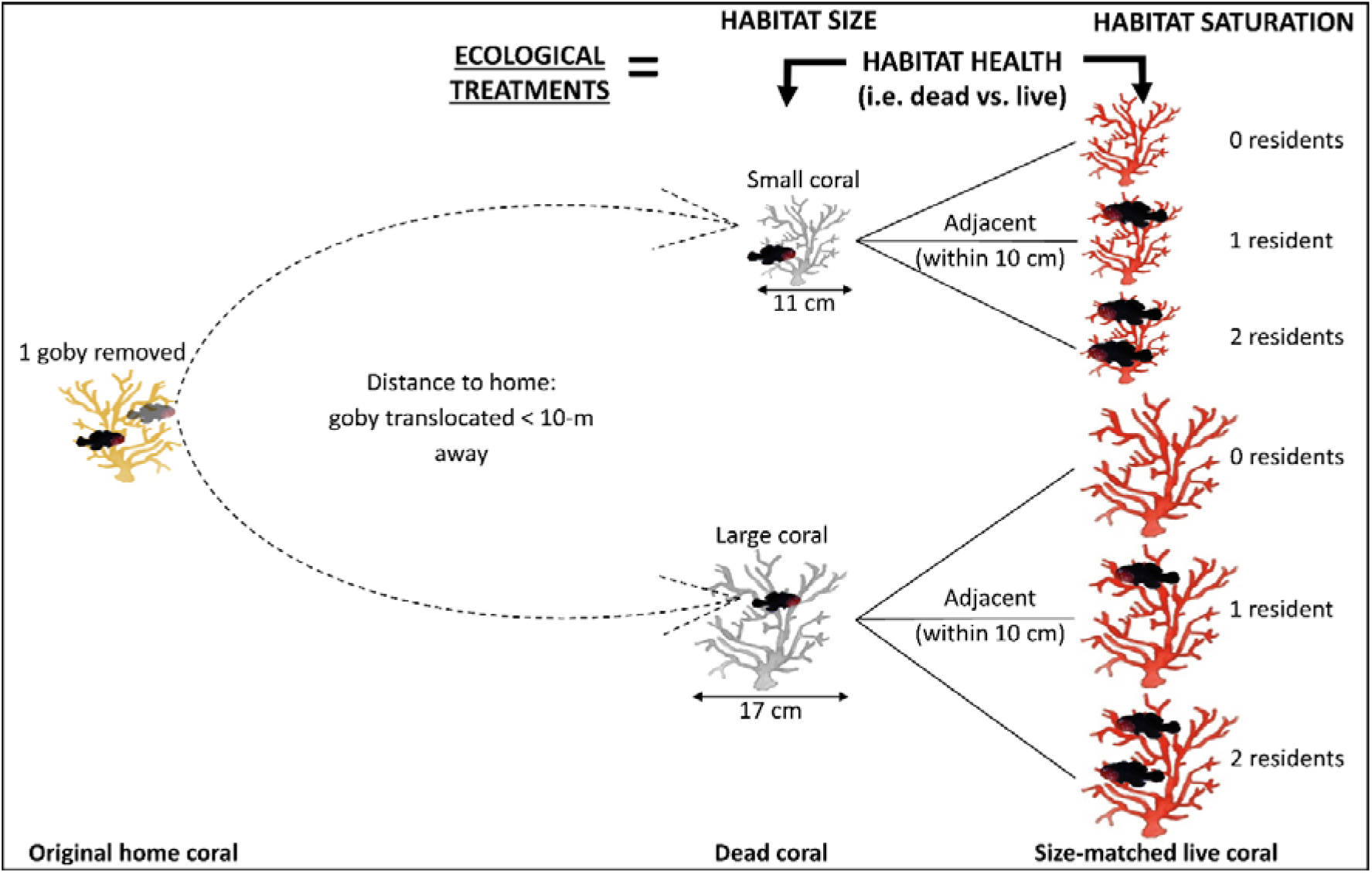
Experimental design: a focal goby was translocated into a dead coral adjacent to an unfamiliar live coral of similar size to offer two habitat health options: dead coral (grey) vs. live coral (red). Six treatment combinations were used to account for two habitat sizes and three habitat saturation levels.

Since the dead coral was adjacent to the live coral, this gave the focal goby the choice of a dead or live coral (thereby examining the effect of habitat health). To simultaneously assess effects of habitat size, the dead and live corals were size-matched in each trial (small or large, Fig 1). In addition, to investigate the role of habitat saturation, treatments were carried out using both small and large coral sizes under three levels of habitat saturation (Fig 1): (i) no residents, (ii) one bigger conspecific, or (iii) two bigger conspecifics in the live coral. Accordingly, a total of six treatment combinations were trialled: three levels of habitat saturation for two levels of habitat size (Fig 1). Ten trials were completed per treatment combination, totalling sixty trials. For each trial, a different focal fish and live coral were used.

To assess where the focal goby moved and whether any movement decisions were based on the level of saturation of neighbouring corals in the study plot, we surveyed all *Acropora* corals larger than 7-cm in diameter [26] within a 10-m radius from the dead coral in each trial. Additional covariables were recorded and accounted for in data analysis (see Suppl Methods).

### Data analysis

The effect of habitat health (live or dead) on the final location of focal gobies was compared using a chi-squared goodness-of-fit test with the null hypothesis that gobies would equally prefer dead and live corals. The effects of the six treatment combinations on the final location of the focal goby (i.e., in dead coral, in live coral, goby not located, returned to home coral) were compared using multinomial logistic regression models. Both habitat size (small or large) and habitat saturation (0, 1, or 2 residents) were included as fixed factors along with the following covariables: distance to home coral, number of gobies in home coral, proportion of uninhabited corals within 10-m radius, and average group size of conspecifics in inhabited corals within 10-m radius. Recruits (distinguished from other life stages by distinct colour and markings, Hing et al. 2018) in the home coral were not included in analysis, because recruits often move between corals before settlement (Froehlich pers. obs. & [16]). The effect of movement costs on the probability of finding the focal goby (would be located [moved successfully] or no longer located [moved unsuccessfully]) was compared using a chi-squared goodness-of-fit test. Data analysis was completed in RStudio [27] with R v4.0.1 [28] packages: VGAM [29], car [30], tidyverse [31], and rcompanion [32].

## Results

We completed 24 trials in 2018 and 36 trials in 2019 (total = 60 trials). Movement decisions of focal gobies were not dependent on habitat health (p < 0.001, see Suppl Tab 1 for all statistical outputs, Fig 2), size (p = 0.93, Fig 2), or saturation (p = 0.88, Fig 2). None of the measured covariables were related to movement decisions (p > 0.12). Surprisingly, most focal gobies (93%, n = 56) did not remain in the dead coral or move to the adjacent live coral. Only one goby stayed in the dead coral and three moved into the live coral (Fig 2). Instead, 55% of focal gobies (n = 33) were located back in their home coral, which was up to 10-m away (Fig 3a). Gobies that returned home travelled between 0.6 to 9 m (Fig 3b). While most returned home within 1 day, some took up to 7 days (Suppl Fig 3). The remaining 38% of gobies (n = 23) could not be located anywhere in the dead coral, live coral, home coral, or in any of the corals within a 10-m radius. Overall, 33 individuals were located and 23 individuals were not located, despite thorough searches, suggesting that they did not survive and faced high costs of movement (p = 0.18).

**Fig 2.**
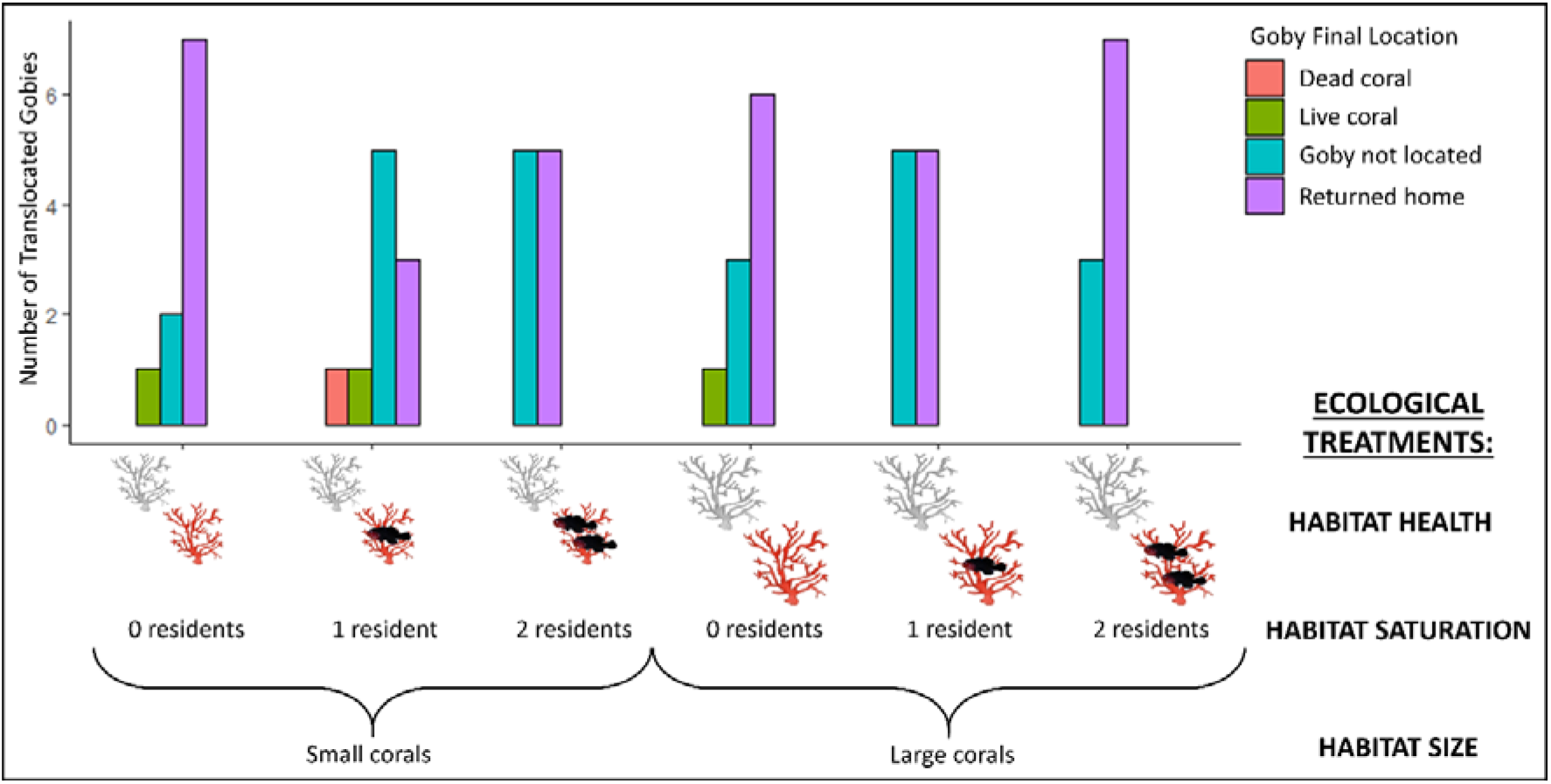
Frequency of gobies’ final location in relation to habitat health (live/red coral or dead/grey coral), saturation and size.

**Fig 3.**
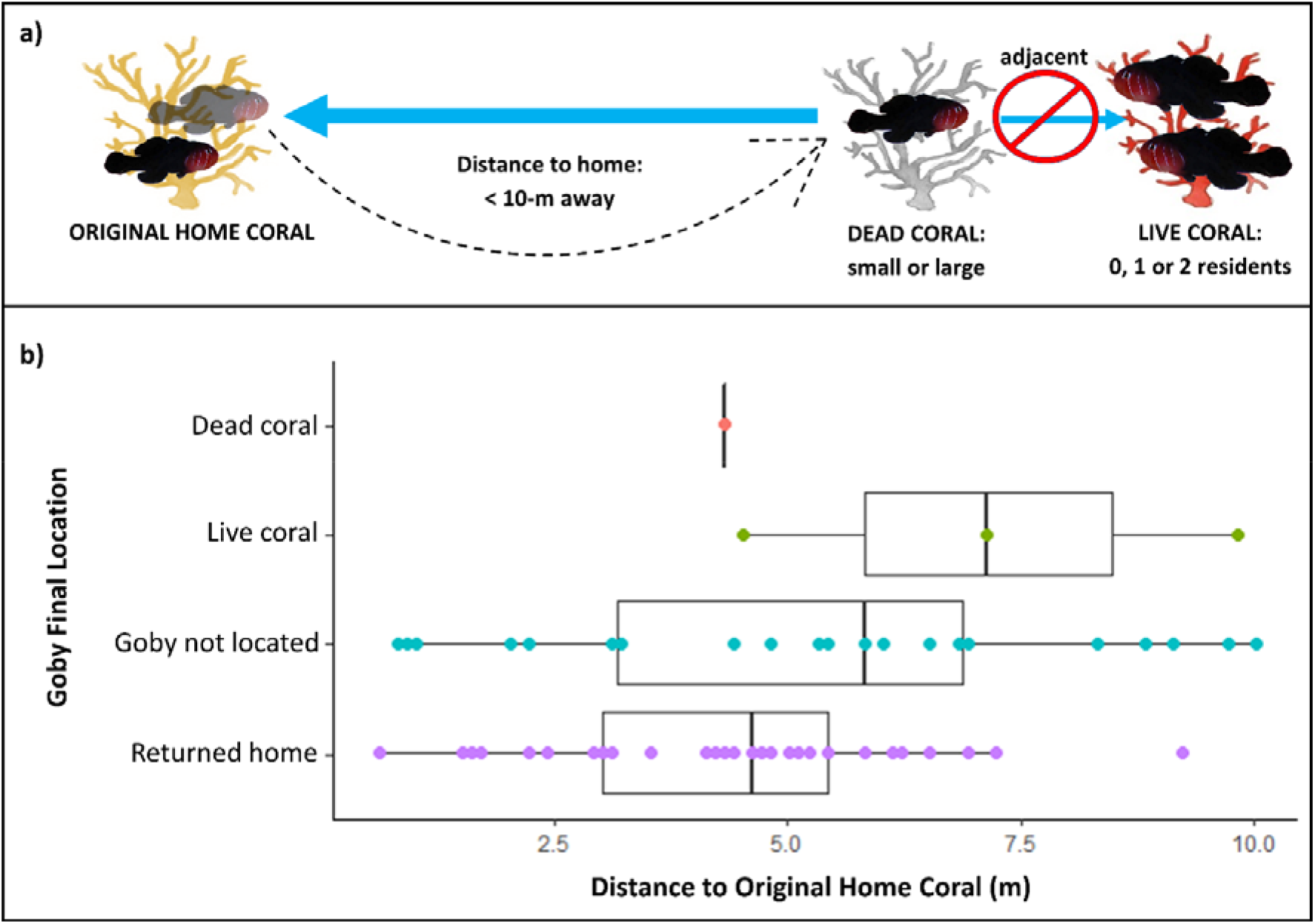
**a** Most common outcome for coral gobies that were translocated into a dead coral away from their home coral (beige). Black dashed arrow represents translocation, blue arrows represent expected outcomes and the red circle crosses out the least popular outcome. **b** Final location of focal gobies in relation to the distance to travel and return to their home coral.

## Discussion

By experimentally manipulating three ecological factors (habitat health, size, saturation), we simultaneously tested multiple components of two hypotheses of sociality: ecological constraints (costs of movement and habitat saturation) and benefits of philopatry (habitat health and size). Surprisingly, when these small-bodied fish were translocated up to 10-m away, they preferentially returned to their home coral instead of moving into an alternative live coral nearby (within 10 cm). This preference occurred despite high apparent costs of movement (38% chance of mortality). In contrast, movement decisions were not related to habitat health, size, and saturation, contradicting the hypothesized role of ecological factors on movement. Instead, these findings highlight an unsung role of habitat familiarity and benefits of homing in movement decisions and hence the maintenance of sociality in this social fish.

For other social reef fishes, previous studies have demonstrated that habitat factors influence the movement decisions of individuals, thereby promoting sociality [12,14]. Numerous studies found positive correlations between habitat size and group size [16,24,33–36], demonstrating the important role of habitat in determining levels of sociality. In addition, habitat saturation influences dispersal and grouping decisions in the coral goby *Paragobiodon xanthosoma* [14] whereby individuals preferentially move to adjacent corals of low saturation (low risk of movement). Furthermore, since coral gobies and damselfishes only inhabit relatively healthy corals and leave highly degraded and dead corals [23,37–39], we expected coral health to influence movement decisions. However, the current study demonstrated that none of these habitat factors (size, saturation and health) influenced the movement of *G. quinquestrigatus*; instead, gobies remarkably returned home even though i) they were often reinstated as nonbreeding subordinates at home, ii) there were opportunities to breed immediately in nearby corals that were healthy, large, and had low saturation, and iii) there were high costs of returning home due to the long distances and risks of predation.

Why do *G. quinquestrigatus* individuals face high costs of movement and return home when other social reef fish species, *P. xanthosoma* and *Amphiprion percula*, prefer instead to join alternative groups [12,14]? Homing ability has already been demonstrated in *G. histrio* [40], other cryptobenthic and reef fishes [41,42], suggesting broader benefits of homing. However, the anemonefish *A. percula* only homed when distances to travel home were small (0.5 m) and never when ecological constraints were heightened and travel distances reached 5 m [12]. Interestingly, even though *G. quinquestrigatus* are at least one third smaller than anemonefish, they preferred to home despite longer distances (up to 10 m) and high costs of movement (estimated 38% mortality). Perhaps *G. quinquestrigatus* home due to the benefits of associating with familiar conspecifics, like in social damselfishes [43]. A well-established social hierarchy [12,44] means avoiding costs of re-establishing dominance, like immediate eviction and possible mortality from enhanced aggression by unfamiliar residents [18,45]. Importantly, since gobies in our study returned home even if they were the only one residing in that coral, there may be benefits of returning to a familiar host habitat, as seen in cardinalfishes [46]. Cardinalfishes move hundreds of meters daily and return to the same host, but host fidelity is more important than mate fidelity because new mates are common [46]. Gobies on the other hand, may move temporarily between corals as juveniles, but eventually select a particular host and never leave that coral [47]. This suggests that certain aspects of their particular coral habitat may enhance their fitness [22]. Thus, choosing an alternative host could be less advantageous than attempting to return to their familiar home coral.

Our results demonstrate that coral gobies are clearly specialized, not only to a particular type of habitat but also to specific habitats that they are familiar with. Such specificity might prove disadvantageous under conditions of rapid habitat degradation, particularly due to cyclones and bleaching [16,21,26], because maintaining plasticity in habitat utilization would enable these fish to reside in any habitat available following environmental disturbances [39]. Unlike other social fishes, however, *G. quinquestrigatus* opted to pay high costs of movement by returning to familiar corals rather than adopting other suitable corals nearby. Such interspecific differences may disproportionally alter the maintenance of sociality among species as their habitats are degrading at alarming rates. Since our study site was located on a relatively pristine reef system, perhaps the enhanced movement frequency of gobies reflects the overall reef condition. Hence, focal individuals may only restrict movements and adopt alternative habitat if their reef system is degraded. Further research investigating whether degrees of disturbance affect movement and grouping decisions would be important for predicting the impacts of environmental change on social species.

## Conclusions

While habitat factors are thought to play an important role in sociality, here we show that habitat saturation, size and health do not influence the use of alternative hosts by coral gobies when their home habitats are still viable. Instead of forming new groups or inhabiting alternative corals of high quality, this social fish opts to swim long distances to return to their familiar home coral. These findings suggest that habitat, mate and/or social group familiarity drives homing behaviour in coral gobies, which in turn is likely to influence the formation and maintenance of their social groups. Our findings therefore question how widely applicable the findings of pre-existing studies are on other social fishes. Future changes to reef environments due to climate change will likely alter the trade-offs of movement as their habitat becomes more fragmented and truncated, which raises doubts about the maintenance of sociality and persistence of populations under future conditions.

## Acknowledgements

We are grateful to the local owners of the reefs: the Kilu and Tamare communities of Papua New Guinea. We thank Theresa Rueger, Nelson and Jerry Sikatura, Somei Jonda, Peter and Jane Miller, the Mahonia Na Dari Research and Conservation Centre, and the Walindi Plantation Resort for field assistance and site visitation.

## Funding

The project was funded by a Sea World Research and Rescue Foundation grant to MW, and funding initiatives from the Centre for Sustainable Ecosystem Solutions at the University of Wollongong (UOW) to CF and MW. CF was supported by a University Postgraduate Award, and SH by an Australian Government Research Training Program Scholarship, both administered through UOW.

### Animal Ethics

University of Wollongong AE1404 and AE1725. Papua New Guinea Research Visa Permit AA654347.

## Data Availability

All data and statistical coding are available at the Knowledge Network for Biocomplexity repository with identifier: *doi:10.5063/D21W00*.

## Notes

https://doi.org/10.5063/D21W00

### Competing Interest Statement

The authors have declared no competing interest.

