## Supplemental Material for "Habitat size, health and saturation do not alter movement decisions or the preference for familiarity in a social coral-reef fish"

### <u>Supplementary Material for Article:</u>

Supplementary Methods:

We located sufficient numbers of live corals containing one or two residents, so we did not need to manipulate group sizes for habitat saturation treatments of one or two gobies. However, there were not enough empty live corals for the treatment with no residents (Fig 1), so we removed all residents from live corals to simulate the lowest saturation level. Removed residents were housed at the Mahonia Na Dari Research and Conservation Centre in a 15l bucket with an air stone, given pellet food twice a day, and received a 25% water change daily. Residents were returned to their home corals 48hrs later.

Before each trial, the focal goby was removed from its home coral and given 5 min to recover in a seawater-filled bag underwater. To start a trial, the focal goby was translocated into the dead coral that was covered in a net to allow the fish to settle in the dead coral without potentially darting off (Suppl Fig 1c). The goby was given a further 5 min to settle before the net was removed. Initially, the focal gobies were observed for 30 min by a scuba diver *in situ* and recorded using a video camera (GoPro 5) for the first 24 trials. However, during these initial observations we noticed that focal gobies rarely moved from the dead coral so subsequent trials only included a 5-min observation by a scuba diver once the net was removed. In the following days, to confirm whether and where focal gobies had moved, the dead coral, adjacent live coral, and home coral were checked daily for two weeks. From these observations, it became apparent that gobies not located within the first week were never located, hence thereafter experimental plots were revisited daily for up to 7 days. Since most gobies moved within the first 24hrs (Suppl Fig 2), if a focal goby stayed in either the dead coral or the unfamiliar live coral for 48hrs, their choice was recorded and the goby was then returned to its home coral.

During our experiments, the following covariables were also recorded on conclusion of each trial: year of experiment, reef, depth, focal goby standard length (mm), distance to its home coral (cm), number of subadult/adult fish in its home coral, species of home coral (Suppl Fig 2), avg. diameter of home coral (as per Kuwamura et al. 1994), species of unfamiliar live coral (Suppl Fig 2), avg. diameter of unfamiliar live coral, number of corals in 10-m radius, avg. goby group size for corals in 10-m radius, proportion of empty corals in 10-m radius, avg. and group size of conspecifics in occupied corals in 10-m radius (see Suppl Tab 1).

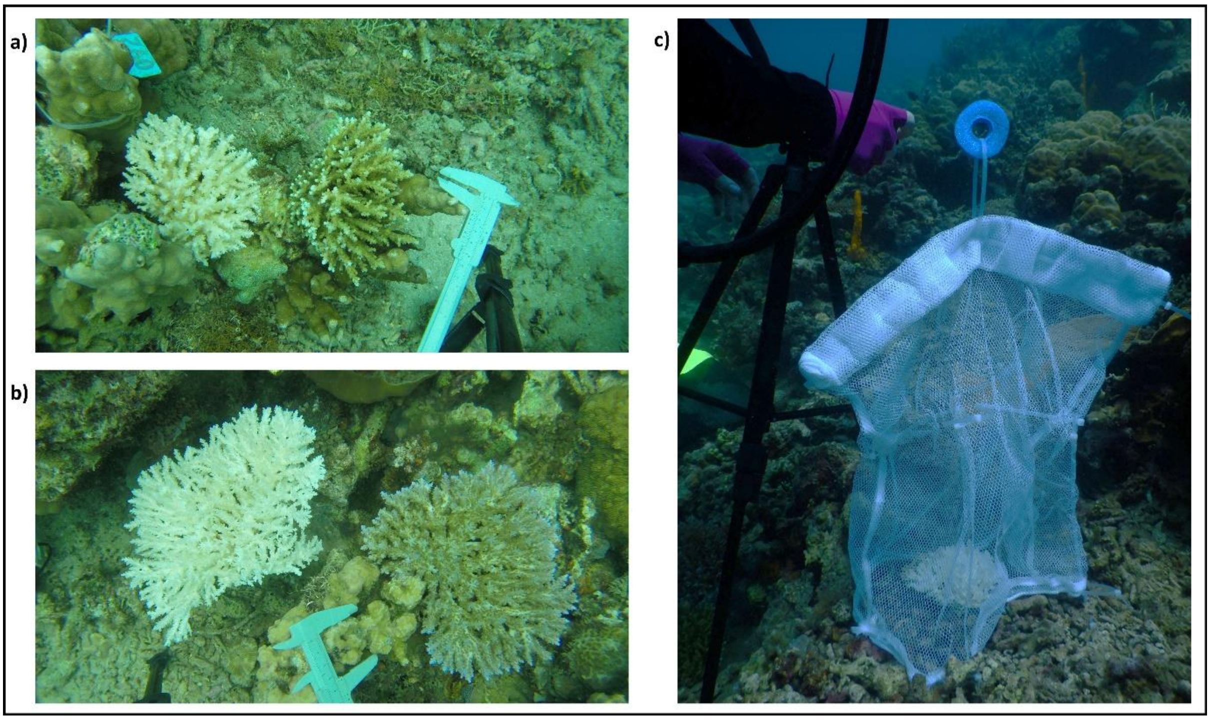


Supplementary Figure 1. The dead coral (white) and unfamiliar live coral were size matched for: **a** small corals and **b** large corals. **c** A net was placed around the dead coral for 5 min at the start of each trial and subsequently removed.

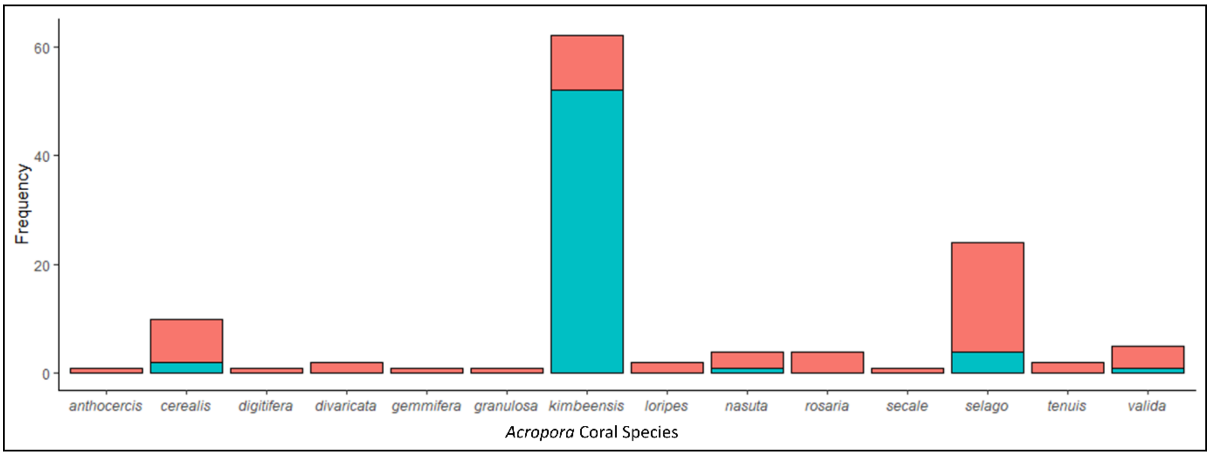


Supplementary Figure 2. Frequency of *Acropora* species that were used in the experiment for the home coral (red) and the unfamiliar live coral (blue).

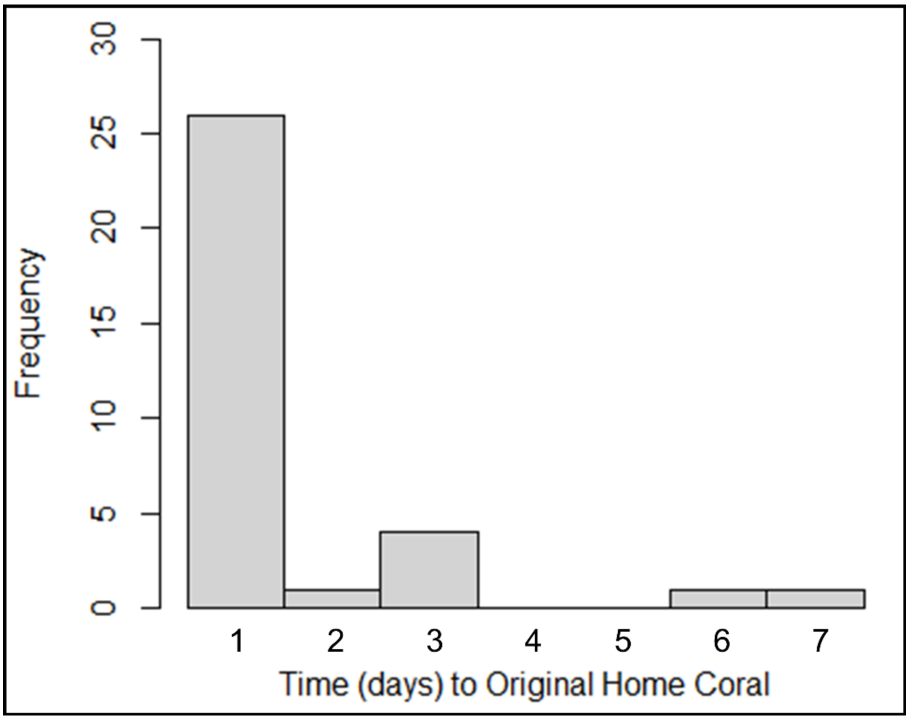

Supplementary Figure 3. Number of days that focal gobies took to return home successfully.

Supplementary Table 1. Statistical outputs of all analyses for the movement decisions of coral dwelling gobies *Gobiodon quinquestrigatus* in relation to covariables and treatments. Additional covariables not discussed in detail in the manuscript are also included here. Habitat size means coral size, and habitat saturation means the number of gobies in the unfamiliar live coral. “Avg.” means average; “Prop.” means proportion; “within 10-m” refers to the survey completed in a 10-m radius around each experimental setup; "N/A" means not applicable.

| Response variable | Predictor variable | Factor Type | Model Test | df | $\chi^2$ value | p-value | pseudo R-squared Statistic | Test Value |
|---|---|---|---|---|---|---|---|---|
| focal goby final location | habitat health | N/A | chi-squared goodness-of-fit test | 3 | 48.533 | < 0.0001 | N/A | N/A |
| if goby was located or not located | cost of movement | N/A | chi-squared goodness-of-fit test | 1 | 1.7857 | 0.1814 | N/A | N/A |
| focal goby final location | habitat size | fixed | multinomial | 5 | 1.3730 | 0.9272 | McFadden | 0.1132 |
| | habitat saturation | fixed | logistic | 8 | 3.7300 | 0.8806 | Cox and Snell (ML) | 0.1870 |
| | interaction: habitat size x saturation | fixed | regression model | 6 | 1.0330 | 0.9843 | Nagelkerke (Cragg and Uhler) | 0.2228 |
| | model likelihood ratio test | | | 15 | 12.4230 | 0.6467 | | |
| focal goby final location | distance to intruder home coral | covariable | multinomial | 3 | 1.9815 | 0.5763 | McFadden | 0.3808 |
| | number of fish in intruder home coral | covariable | logistic | 3 | 0.7351 | 0.8649 | Cox and Snell (ML) | 0.5016 |
| | prop. uninhabited corals within 10-m | covariable | regression | 3 | 1.0015 | 0.8009 | Nagelkerke | 0.5976 |
| | avg. group size of conspecifics in corals within 10-m | covariable | model | 3 | 0.7680 | 0.8571 | (Cragg and Uhler) | |
| | habitat size | fixed | | 5 | 2.4247 | 0.7878 | | |
| | habitat saturation | fixed | | 8 | 3.8077 | 0.8740 | | |
| | interaction: habitat size x saturation | fixed | | 6 | 2.0385 | 0.9161 | | |
| | model likelihood ratio test | | | 27 | 41.7860 | 0.0346 | | |
| focal goby final location | distance to intruder home coral | covariable | multinomial | 3 | 4.0416 | 0.2570 | McFadden | 0.1885 |
| | habitat size | fixed | logistic | 4 | 0.6953 | 0.9519 | Cox and Snell (ML) | 0.2915 |
| | habitat saturation | fixed | regression | 8 | 4.1305 | 0.8452 | Nagelkerke | 0.3473 |
| | interaction: habitat size x saturation | fixed | model | 6 | 1.1887 | 0.9774 | (Cragg and Uhler) | |
| | model likelihood ratio test | | | 18 | 20.6800 | 0.2959 | | |
| focal goby final location | number of fish in intruder home coral | covariable | multinomial | 3 | 0.4837 | 0.9225 | McFadden | 0.2012 |
| | habitat size | fixed | logistic | 4 | 0.9593 | 0.9159 | Cox and Snell (ML) | 0.3079 |
| | habitat saturation | fixed | regression | 8 | 3.9462 | 0.8619 | Nagelkerke | 0.3668 |
| | interaction: habitat size x saturation | fixed | model | 6 | 1.1645 | 0.9786 | (Cragg and Uhler) | |
| | model likelihood ratio test | | | 18 | 22.0770 | 0.2286 | | |
| focal goby final location | unfamiliar live coral average diameter | covariable | multinomial | 3 | 0.0769 | 0.9945 | McFadden | 0.1139 |
| | habitat size | fixed | logistic | 5 | 1.2397 | 0.9410 | Cox and Snell (ML) | 0.1881 |
| | habitat saturation | fixed | regression | 8 | 3.6473 | 0.8875 | Nagelkerke | 0.2240 |
| | interaction: habitat size x saturation | fixed | model | 6 | 1.0695 | 0.9828 | (Cragg and Uhler) | |
| | model likelihood ratio test | | | 18 | 12.5000 | 0.8204 | | |
| focal goby final location | intruder home coral average diameter | covariable | multinomial | 3 | 2.4219 | 0.4896 | McFadden | 0.1460 |
| | habitat size | fixed | logistic | 4 | 1.5723 | 0.8138 | Cox and Snell (ML) | 0.2344 |
| | habitat saturation | fixed | regression | 8 | 3.7882 | 0.8757 | Nagelkerke | 0.2792 |
| | interaction: habitat size x saturation | fixed | model | 6 | 0.6783 | 0.9949 | (Cragg and Uhler) | |
| | model likelihood ratio test | | | 18 | 16.0220 | 0.5910 | | |
| focal goby final location | number of corals within 10-m | covariable | multinomial | 3 | 3.2425 | 0.3557 | McFadden | 0.1463 |
| | habitat size | fixed | logistic | 5 | 2.3435 | 0.7999 | Cox and Snell (ML) | 0.2348 |
| | habitat saturation | fixed | regression | 8 | 4.2744 | 0.8316 | Nagelkerke | 0.2797 |
| | interaction: habitat size x saturation | fixed | model | 6 | 1.9181 | 0.9271 | (Cragg and Uhler) | |
| | model likelihood ratio test | | | 18 | 16.0550 | 0.5887 | | |
| focal goby final location | reef sampled | covariable | multinomial | 9 | 2.5301 | 0.9801 | McFadden | 0.1668 |
| | habitat size | fixed | logistic | 5 | 1.9122 | 0.8612 | Cox and Snell (ML) | 0.2629 |
| | habitat saturation | fixed | regression | 8 | 3.7378 | 0.8800 | Nagelkerke | 0.3131 |
| | interaction: habitat size x saturation | fixed | model | 6 | 1.2359 | 0.9751 | (Cragg and Uhler) | |
| | model likelihood ratio test | | | 24 | 18.2980 | 0.7883 | | |

Supplementary Table 1 continued.

| Response variable | Predictor variable | Factor Type | Model Test | df | $\chi^2$ value | p-value | pseudo R-squared Statistic | Test Value |
|---|---|---|---|---|---|---|---|---|
| focal goby | avg. group size in corals within 10-m | covariable | multinomial | 3 | 2.3528 | 0.5025 | McFadden | 0.1382 |
| final location | habitat size | fixed | logistic | 5 | 1.5734 | 0.9044 | Cox and Snell (ML) | 0.2233 |
| | habitat saturation | fixed | regression | 8 | 3.9323 | 0.8632 | Nagelkerke | 0.2660 |
| | interaction: habitat size x saturation | fixed | model | 6 | 1.1439 | 0.9796 | (Cragg and Uhler) | |
| | model likelihood ratio test | | | 18 | 15.1610 | 0.6509 | | |
| focal goby | prop. corals within 10-m with conspecifics | covariable | multinomial | 3 | 5.6503 | 0.1299 | McFadden | 0.2267 |
| final location | habitat size | fixed | logistic | 5 | 2.5984 | 0.7616 | Cox and Snell (ML) | 0.3394 |
| | habitat saturation | fixed | regression | 8 | 4.5473 | 0.8047 | Nagelkerke | 0.4043 |
| | interaction: habitat size x saturation | fixed | model | 6 | 1.1184 | 0.9807 | (Cragg and Uhler) | |
| | model likelihood ratio test | | | 18 | 24.8750 | 0.1284 | | |
| focal goby | avg. group size of conspecifics in corals within 10-m | covariable | multinomial | 3 | 2.8486 | 0.4156 | McFadden | 0.1444 |
| final location | habitat size | fixed | logistic | 5 | 1.7018 | 0.8887 | Cox and Snell (ML) | 0.2320 |
| | habitat saturation | fixed | regression | 8 | 4.1718 | 0.8413 | Nagelkerke | 0.2764 |
| | interaction: habitat size x saturation | fixed | model | 6 | 1.2342 | 0.9752 | (Cragg and Uhler) | |
| | model likelihood ratio test | | | 18 | 15.8410 | 0.6037 | | |
| focal goby | prop. uninhabited corals within 10-m | covariable | multinomial | 3 | 0.9194 | 0.8207 | McFadden | 0.1226 |
| final location | habitat size | fixed | logistic | 5 | 1.3318 | 0.9316 | Cox and Snell (ML) | 0.2008 |
| | habitat saturation | fixed | regression | 8 | 3.6067 | 0.8907 | Nagelkerke | 0.2392 |
| | interaction: habitat size x saturation | fixed | model | 6 | 1.0074 | 0.9853 | (Cragg and Uhler) | |
| | model likelihood ratio test | | | 18 | 13.4490 | 0.7642 | | |
| focal goby | emperimental depth (unfamiliar live coral) | covariable | multinomial | 3 | 2.5315 | 0.4696 | McFadden | 0.1390 |
| final location | habitat size | fixed | logistic | 5 | 1.4310 | 0.9209 | Cox and Snell (ML) | 0.2245 |
| | habitat saturation | fixed | regression | 8 | 3.9675 | 0.8600 | Nagelkerke | 0.2674 |
| | interaction: habitat size x saturation | fixed | model | 6 | 1.0879 | 0.9821 | (Cragg and Uhler) | |
| | model likelihood ratio test | | | 18 | 15.2530 | 0.6445 | | |
| focal goby | year of experiment | covariable | multinomial | 3 | 0.2992 | 0.9602 | McFadden | 0.1229 |
| final location | habitat size | fixed | logistic | 5 | 1.3776 | 0.9267 | Cox and Snell (ML) | 0.2013 |
| | habitat saturation | fixed | regression | 8 | 3.2511 | 0.9176 | Nagelkerke | 0.2398 |
| | interaction: habitat size x saturation | fixed | model | 6 | 0.9905 | 0.9860 | (Cragg and Uhler) | |
| | model likelihood ratio test | | | 18 | 13.4850 | 0.7620 | | |
| focal goby | intruder goby standard length | covariable | multinomial | 3 | 2.6132 | 0.4552 | McFadden | 0.2765 |
| final location | habitat size | fixed | logistic | 5 | 2.5858 | 0.7635 | Cox and Snell (ML) | 0.4240 |
| | habitat saturation | fixed | regression | 8 | 5.1707 | 0.7392 | Nagelkerke | 0.4908 |
| | interaction: habitat size x saturation | fixed | model | 6 | 1.1977 | 0.9770 | (Cragg and Uhler) | |
| | model likelihood ratio test | | | 33 | 30.3440 | 0.6000 | | |
| focal goby | unfamiliar live coral species | covariable | multinomial | 12 | 2.3849 | 0.9985 | McFadden | 0.1727 |
| final location | habitat size | fixed | logistic | 5 | 0.3060 | 0.9975 | Cox and Snell (ML) | 0.2708 |
| | habitat saturation | fixed | regression | 8 | 3.3812 | 0.9082 | Nagelkerke | 0.3227 |
| | interaction: habitat size x saturation | fixed | model | 6 | 0.2565 | 0.9997 | (Cragg and Uhler) | |
| | model likelihood ratio test | | | 27 | 18.9510 | 0.8718 | | |
| focal goby | intruder home coral species | covariable | multinomial | 39 | 2.1569 | 1.0000 | McFadden | 0.3711 |
| final location | habitat size | fixed | logistic | 5 | 1.6304 | 0.8976 | Cox and Snell (ML) | 0.4927 |
| | habitat saturation | fixed | regression | 8 | 4.5299 | 0.8064 | Nagelkerke | 0.5869 |
| | interaction: habitat size x saturation | fixed | model | 6 | 1.6188 | 0.9512 | (Cragg and Uhler) | |
| | model likelihood ratio test | | | 54 | 40.7150 | 0.9092 | | |
